# Utilizing data imbalance to enhance compound-protein interaction prediction models

**DOI:** 10.1101/2024.10.23.619867

**Authors:** Wei Lin, Chi Chung Alan Fung

## Abstract

Identifying potential compounds for target proteins is crucial in drug discovery. Current compound-protein interaction prediction models concentrate on utilizing more complex features to enhance capabilities, but this often incurs substantial computational burdens. Indeed, this issue arises from the limited understanding of data imbalance between proteins and compounds, leading to insufficient optimization of protein encoders. Therefore, we introduce a sequence-based predictor named FilmCPI, designed to utilize data imbalance to learn proteins with their numerous corresponding compounds. FilmCPI consistently outperforms baseline models across diverse datasets and split strategies, and its generalization to unseen proteins becomes more pronounced as the datasets expand. Notably, FilmCPI can be transferred to unseen protein families with sequence-based data from other families, exhibiting its practicability. The effectiveness of FilmCPI is attributed to different optimization speeds for diverse encoders, elucidating optimization imbalance in compound-protein prediction models. Additionally, these advantages of FilmCPI do not depend on increasing parameters, aiming to lighten model design with data imbalance.

## Introduction

Identifying potential drugs for target proteins from numerous candidate compounds is a crucial step in the drug discovery process. Despite significant advancements in compound-protein interaction (CPI) predictors driven by artificial intelligence, ^1^ these models still face a trade-off between effective drug discovery and high computational costs. Traditional sequence-based methods^2–6^ have made strides by incorporating richer molecular features. Even though compound features have evolved from Simplified Molecular-Input Line Entry System (SMILES)^7^ to complex molecular graphs, they still struggle to generalize to unseen proteins^5,8^ owing to the limited information provided by the amino acid sequences. To address this issue, current solutions utilize large pre-trained protein models such as ESM2^9^ or AlphaFold2^10^ to provide richer protein information. ^8,11,12^ However, billions of parameters in ESM2 result in massive memory usage, and AlphaFold2 requires searching in protein structure databases,^13^ leading to substantial time consumption. Consequently, the use of pre-trained protein models enhances the capacities of models, but it also incurs considerable computational costs. Furthermore, the vast drug space,^14^ estimated to be 10^60^, makes these high-cost computational methods impractical for most researchers. Therefore, there is an urgent need for efficient solutions that enhance generalization to unseen proteins.

Perhaps we need to rethink the former notion of adding more complex features to enhance model performance. Actually, the answers may lie in addressing the overlooked issue of data imbalance between proteins and compounds. To be specific, the full BindingDB^15^ dataset contains millions of compounds but only thousands of proteins. According to our analysis exhibited in Fig.2a,c, proteins are significantly larger than compounds in size. These two factors combined complicate the optimization of protein encoder.^16^ Upon thorough analysis, we provide a novel thought: why not focus on learning protein representations alongside their massive corresponding compounds? To validate this idea, we introduce a sequence-based CPI predictor named FilmCPI for experimental purposes. Specifically, we employ compound features that do not require training and thus employ sparse extended-connectivity fingerprints (ECFPs),^17^ which have been widely used for drug discovery.^18^ Then, we adopt the compound representations to scale and shift the protein representations with a feature-wise linear modulation^19^ (Film) module.

We conduct comprehensive experiments on FilmCPI and discover that its advantages are reflected in four aspects. Firstly, FilmCPI consistently outperforms some representative baselines and its variants across diverse datasets and split strategies, indicating its robustness, as shown in Fig.3a and Table1,2. Secondly, as exhibited in Fig.3a, the advantages of FilmCPI to unseen proteins become more pronounced as the datasets expand, implying its scalability. Thirdly, as illustrated in Fig.3b, FilmCPI can effectively utilize CPI data from other protein families to predict interactions in unseen G-protein-coupled receptors (GPCRs) and ion channels, showing its transferability. Finally, although FilmCPI has obtained these advantages, its success does not rely on significantly increasing parameters, as illustrated in Fig.2d. To explore the underlying mechanisms of these advantages, we examine the optimization dynamics of FilmCPI and its variants. Intriguingly, the success of FilmCPI lies in both the minimal optimization for the compound encoder and the greatest optimization for the protein encoder. This optimization imbalance is consistent with our original argument about focusing on optimizing the protein encoder and the choice of ECFPs as the compound inputs. Collectively, these works establish a connection between data imbalance and optimization imbalance, offering valuable insights for future model design.

**Table 1.**
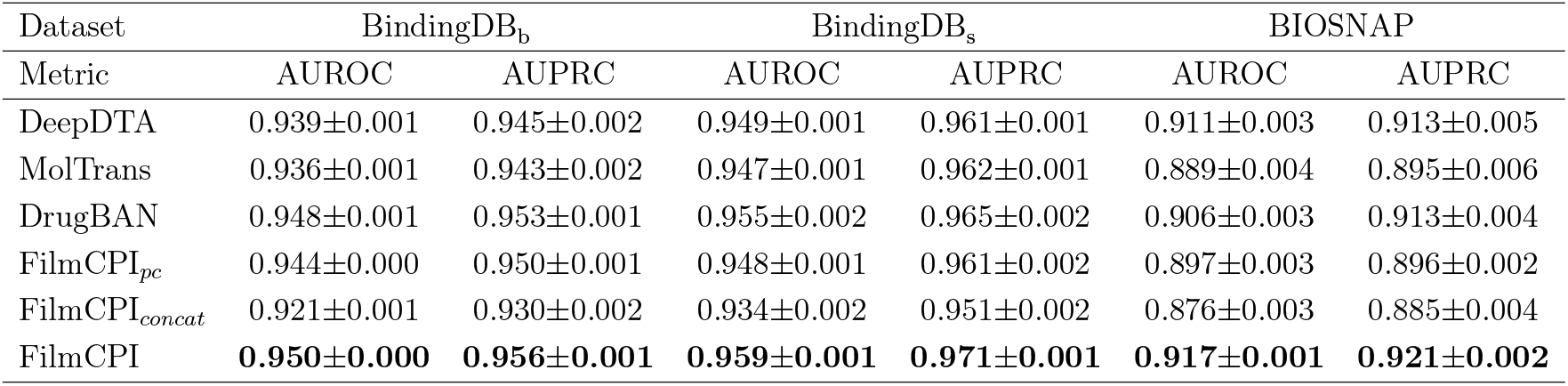
Performance (average AUROC and AUPRC over five random runs) comparison on BindingDB_b_, BindingDB_s_ and BIOSNAP with the random split.

| Dataset | BindingDB<sub>b</sub> |  | BindingDB<sub>s</sub> |  | BIOSNAP |  |
| --- | --- | --- | --- | --- | --- | --- |
| Metric | AUROC | AUPRC | AUROC | AUPRC | AUROC | AUPRC |
| DeepDTA | 0.939±0.001 | 0.945±0.002 | 0.949±0.001 | 0.961±0.001 | 0.911±0.003 | 0.913±0.005 |
| MolTrans | 0.936±0.001 | 0.943±0.002 | 0.947±0.001 | 0.962±0.001 | 0.889±0.004 | 0.895±0.006 |
| DrugBAN | 0.948±0.001 | 0.953±0.001 | 0.955±0.002 | 0.965±0.002 | 0.906±0.003 | 0.913±0.004 |
| FilmCPI<sub>pc</sub> | 0.944±0.000 | 0.950±0.001 | 0.948±0.001 | 0.961±0.002 | 0.897±0.003 | 0.896±0.002 |
| FilmCPI<sub>concat</sub> | 0.921±0.001 | 0.930±0.002 | 0.934±0.002 | 0.951±0.002 | 0.876±0.003 | 0.885±0.004 |
| FilmCPI | **0.950±0.000** | **0.956±0.001** | **0.959±0.001** | **0.971±0.001** | **0.917±0.001** | **0.921±0.002** |

**Table 2.** Performance (average AUROC and AUPRC over five random runs) comparison on BindingDB_b_, BindingDB_s_ and BIOSNAP with the unseen compound split.

| Dataset | BindingDB<sub>b</sub> |  | BindingDB<sub>s</sub> |  | BIOSNAP |  |
| --- | --- | --- | --- | --- | --- | --- |
| Metric | AUROC | AUPRC | AUROC | AUPRC | AUROC | AUPRC |
| DeepDTA | 0.914±0.003 | 0.919±0.004 | 0.914±0.007 | 0.935±0.004 | 0.867±0.007 | 0.877±0.006 |
| MolTrans | 0.908±0.001 | 0.918±0.002 | 0.902±0.007 | 0.926±0.004 | 0.850±0.004 | 0.860±0.005 |
| DrugBAN | 0.932±0.001 | 0.939±0.002 | 0.928±0.008 | 0.945±0.005 | **0.877±0.005** | 0.883±0.006 |
| FilmCPI<sub>pc</sub> | 0.931±0.001 | 0.937±0.002 | 0.927±0.005 | 0.945±0.003 | 0.854±0.007 | 0.860±0.004 |
| FilmCPI<sub>concat</sub> | 0.897±0.003 | 0.908±0.004 | 0.897±0.005 | 0.923±0.004 | 0.841±0.004 | 0.852±0.005 |
| FilmCPI | **0.935±0.003** | **0.942±0.003** | **0.934±0.009** | **0.952±0.005** | **0.877±0.006** | **0.888±0.004** |

## Results

### Problem definition

Given a compound-protein pair, the task aims to determine whether there exists an interaction between the protein and the compound. In most datasets, SMILES is used to represent compounds. For any compound **c**_*i*_, we convert its SMILES into the corresponding ECFP 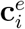 in this work. For any protein **p**_*j*_, we only utilize its amino acids sequences 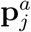.

The objective of CPI models is to learn the mapping function, 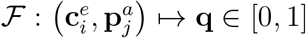, from a compound-protein pair to the interaction probability score.

### FilmCPI framework

The framework of the proposed FilmCPI is exhibited in Fig.1a. Given a compound–protein pair as input, the model first adopts separate, fully connected layers and 1D convolutional neural network (CNN) modules to obtain the representations of molecular fingerprints and protein sequences, respectively. Next, a Film module is utilized to conditionally normalize the protein representations with the corresponding compound representations. Finally, a fully connected classification layer is employed to predict the interaction score between the input compound-protein pair.

**Figure 1.**
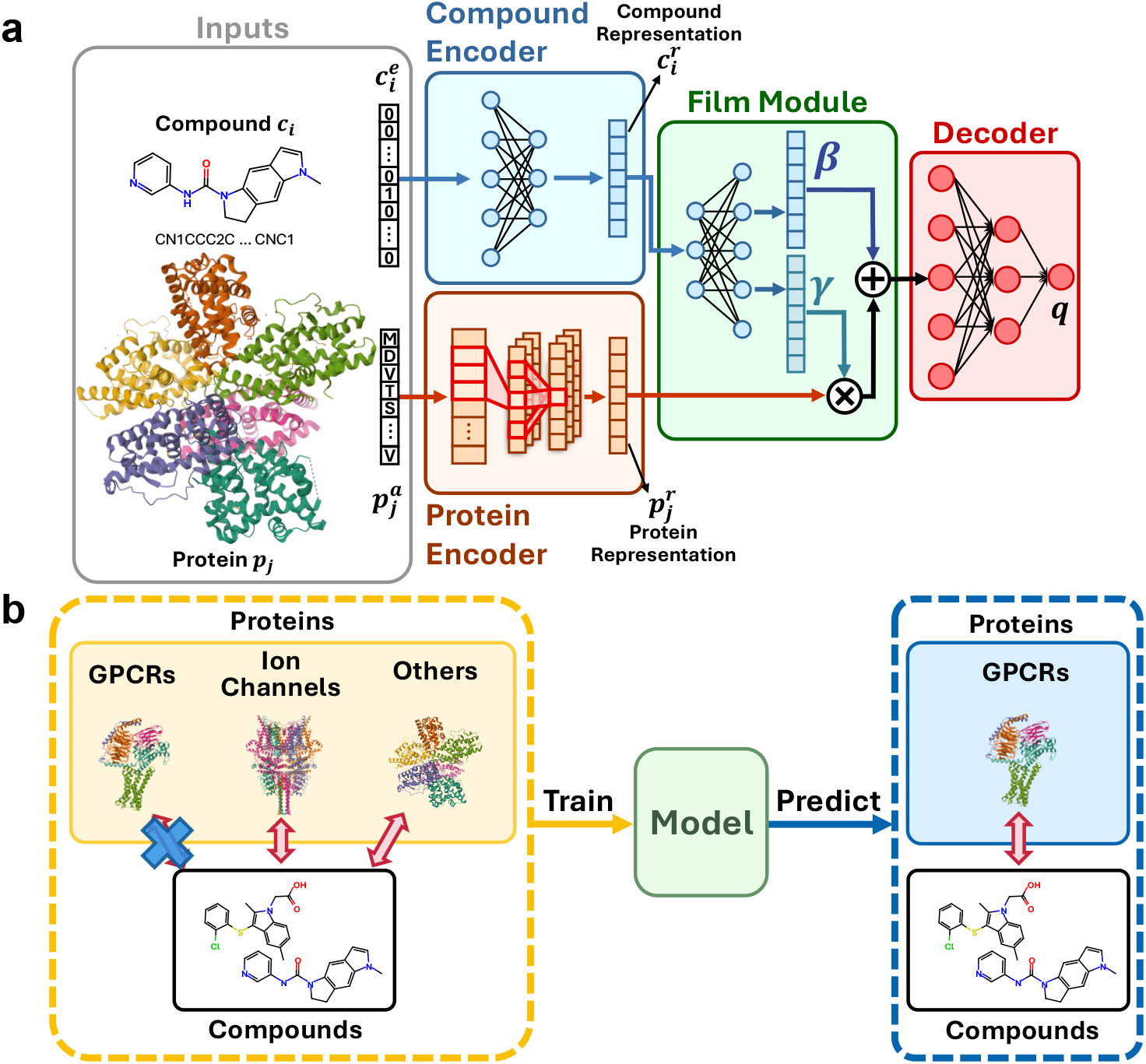
Illustration of FilmCPI and cross-family evaluation. **a**, The input molecular fingerprints 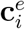 and protein amino acid sequences 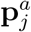 are encoded by fully-connected layers and convolutional neural networks to obtain their representations 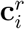 and 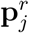, respectively. The representations are integrated with a feature-wise linear modulation module to obtain the interaction probability score **q. b**, We take GPCRs as examples. In the setting of cross-family split, the model is trained with CPI data excluding GPCRs, aiming to predict the CPI data on GPCRs.

### Evaluation strategies and metrics

To comprehensively evaluate the values of FilmCPI, we conduct classification tasks in two phases: traditional evaluation and cross-family evaluation.

Traditional evaluation denotes that we split datasets with three traditional types, namely random split, unseen compound split and unseen protein split, which have been widely used in previous works.^5,6^ Then, we select three public datasets of varying sizes, namely BindingDB_b_, BindingDB_s_, and BIOSNAP,^20^ as the benchmarks. BIOSNAP is created by the previous work.^6^ Both BindingDB_b_ and BindingDB_s_ are extracted from the full BindingDB^15^ dataset, with BindingDB_b_ being the bigger set but BindingDB_s_ being the smaller one. The only difference is the threshold^21^ for different datasets. In this setting, we split the datasets into train/validation/test sets with a ratio of 7:1:2, respectively.

Membrane proteins have an important role in the human body,^22^ making it a crucial component in drug discovery. However, studying the structures of membrane proteins poses significant challenges due to their partially hydrophobic surfaces, flexibility and inherent instability.^23^ In addition, the sequence-based CPI data for some specific membrane protein families only takes a small portion of total CPI data, as exhibited in Supplementary Table1,2. Therefore, we propose a new evaluation method named cross-family evaluation to examine whether these models can be transferred to smaller protein families using CPI data from other families, shown in Fig.1b. We select two important membrane protein families, namely GPCRs^24^ and ion channels,^25^ from BindingDB_b_, which is the largest dataset in this study. We first take interactions from GPCRs or ion channels, respectively. Then, the remaining interactions are divided into train and validation sets with a ratio of 8:2 through the random split, aiming to fully utilize CPI data from other families.

For the above situations, we employ two evaluation metrics, namely the area under the receiver operating characteristic curve (AUROC) and the area under the precision-recall curve (AUPRC), to evaluate model classification performance.

### Traditional performance comparison

In this part, we compare FilmCPI with three representative baselines: DeepDTA, ^2^ MolTrans,^6^ and DrugBAN.^5^ These models have distinct compound features in contrast to FilmCPI. Besides, we introduce two variants of FilmCPI, namely FilmCPI_pc_ and FilmCPI_concat_, as baselines. These variants exhibit minor differences in the fusion operation. As such, they serve to ascertain whether the advancements observed in FilmCPI are dependent on the overall architectural design rather than stemming solely from simple modifications to certain modules. Furthermore, based on the concepts underlying our model design, we propose the first argument: if each protein has more corresponding compounds in the dataset, the model will obtain enhanced generalization to unseen proteins.

The results of the three traditional splits are exhibited in Fig.3a, and Table1,2. From Fig.3a, we observe that the advantages of FilmCPI for unseen proteins increase with the expansion of datasets, supporting our first argument and exhibiting its scalability. In contrast, the trend is reversed if the positions of protein and compound are swapped in FilmCPI, which is FilmCPI_pc_. The phenomenon of FilmCPI_pc_ indirectly validates the first argument.

According to Table1,2, FilmCPI also achieves the best outcomes in both random split and unseen compound split. Unlike FilmCPI, the other models struggle to maintain robust performance across diverse evaluation settings. For instance, DeepDTA is the second-best model for unseen proteins, but its performance declines in the unseen compound split and random split with the expansion of datasets. On the other hand, DrugBAN shows competitive results in random split and unseen compound split, but its generalization to unseen proteins is poor when the dataset is small. These observations further validate the robustness of the FilmCPI. However, they also raise intriguing questions about the underlying mechanism of these models, which will be explored and discussed in the following sections. Numerical results of the unseen protein split are shown in Supplementary Table 3.

### Cross-family performance comparison

Since the available CPI data is limited for certain protein families, it is crucial to examine whether FilmCPI has the ability to transfer its capabilities to unseen protein families, leveraging data from other families. According to the above analysis, we select two significant membrane protein families, GPCRs and ion channels, and compare the performance of FilmCPI against three baseline models.

As shown in Fig.3b, FilmCPI demonstrates outstanding and robust performance compared to other baselines. Specifically, it achieves 11.4% and 14.7% improvements over random guess on AUROC for GPCRs and ion channels, respectively. Besides, the other models exhibit data points that fall below the random guess level on AUROC for these two membrane protein families, a pitfall that does not occur with the FilmCPI model. Notably, GPCRs, ion channels and other proteins display considerable differences in structure and length, as shown in Fig.1b, Fig.2b. This emphasizes the challenge of this task and further highlights the values of FilmCPI. The numerical results of the cross-family split are shown in Supplementary Table 4.

**Figure 2.**
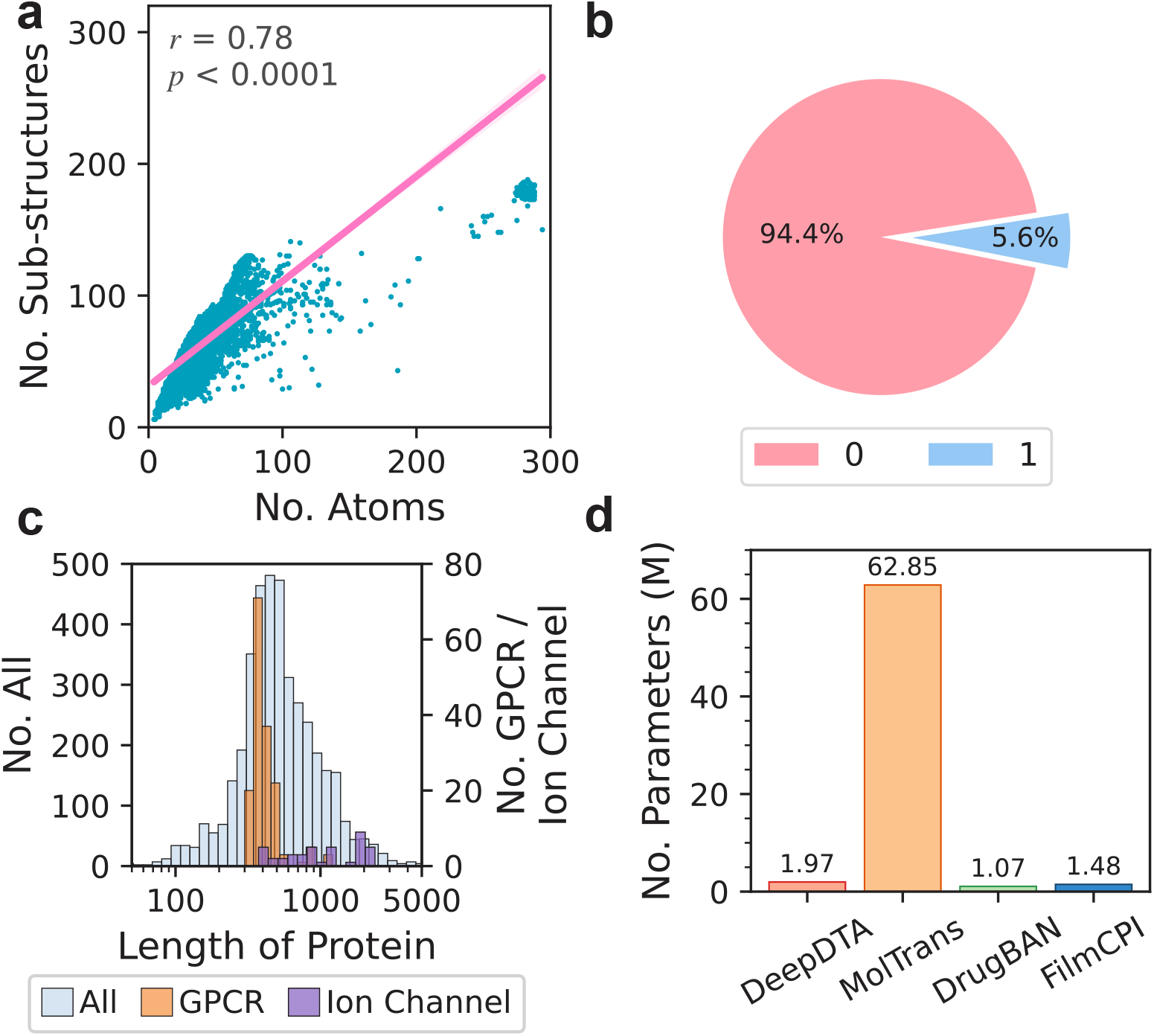
Data analysis of proteins, compounds, and parameters in diverse models. **a**, Correlation between the number of atoms and the number of specific sub-structures in all compounds from BindingDB_b_. **b**, Occupancy of all specific sub-structures in ECFP4 fingerprints, each with a length of 1024, from BindingDB_b_. **c**, Length distributions of proteins, including all proteins, GPCRs, and ion channels in BindingDB_b_. **d**, Number of parameters in sequence-based models used in this work, including DeepDTA, MolTrans, DrugBAN, and FilmCPI.

**Figure 3.**
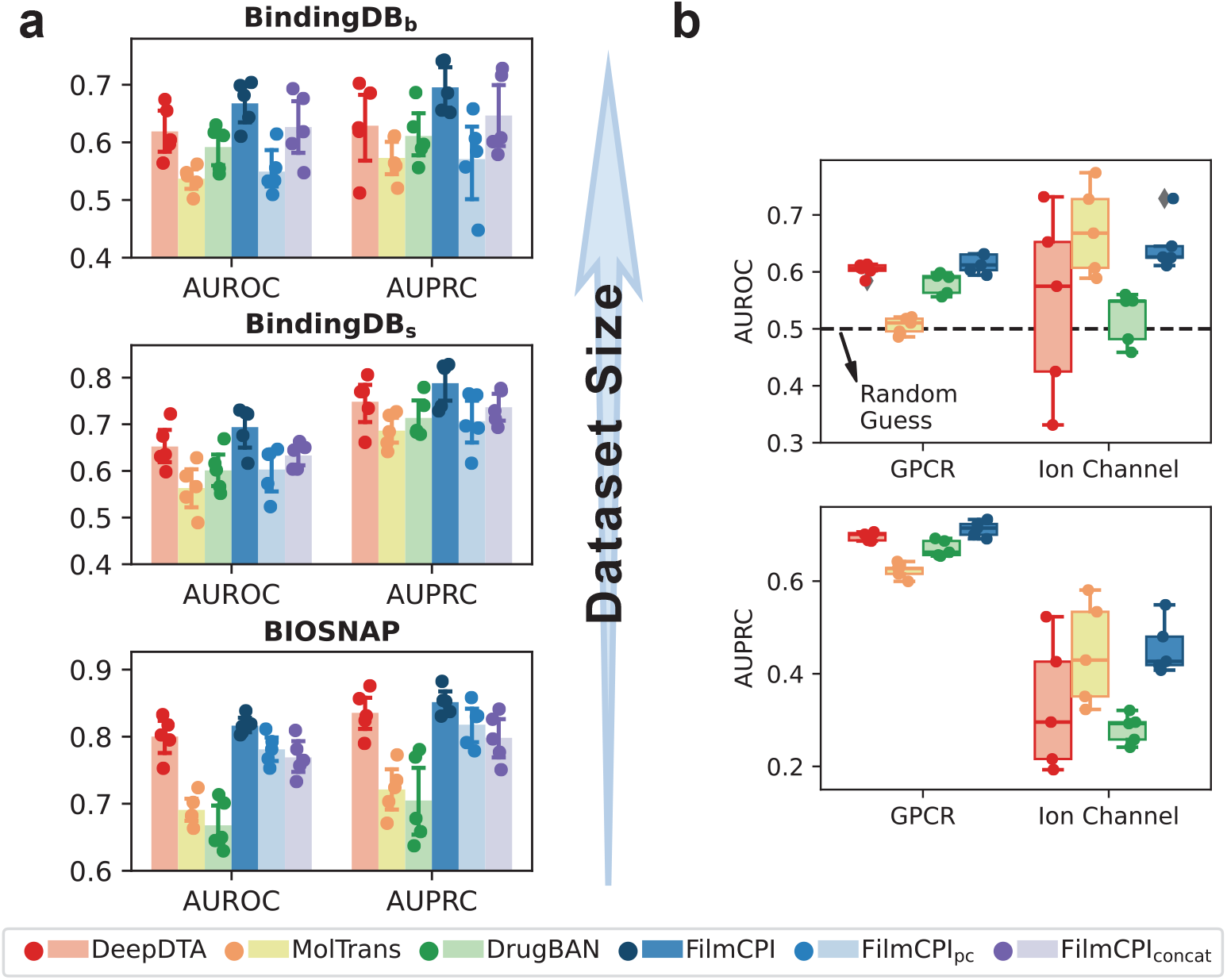
Performance of models to unseen proteins with different split strategies. **a**, Performance of FilmCPI compared with three baselines and two of its variants using the unseen protein split. Upper panel: performance of models in BindingDB_b_. Middle panel: performance of models in BindingDB_s_. Lower panel: performance of models in BIOSNAP. Number of samples: five. **b**, Performance of FilmCPI compared with three baseline models using the cross-family split (GPCRs and ion channels).

### Analysis of optimization dynamics

The phenomena of DeepDTA and DrugBAN, illustrated in the previous section, also happen to FilmCPI_pc_ and FilmCPI_concat_. To be specific, FilmCPI_pc_ performs poorly in the unseen protein split, while FilmCPI_concat_ struggles in both the random split and the unseen compound split. To further investigate the peculiar phenomena exhibited by these models, we train FilmCPI and its variants on BindingDB_b_, excluding the GPCR protein family. In this section, we present the second argument: in terms of optimization dynamics, the optimization should center on the protein encoder rather than the compound encoder.

We start by visualizing the training loss and the test loss of FilmCPI and its variants, as exhibited in Fig.4a. We observe that optimizing FilmCPI_concat_ is more challenging, which explains why previous approaches aimed to enhance the fusion of compound and protein representations. Furthermore, FilmCPI exhibits little improvement over FilmCPI_pc_ according to loss. To understand the optimization dynamics, we visualize the parameter distributions of the encoders, inspired by previous work.^16^ However, the differences across FilmCPI and FilmCPI_pc_ in both encoders are small, as exhibited in Fig.4b,c. Thus, we visualize the variances of the compound representations and the protein representations for unseen GPCRs and seen ion channels as the epoch changes, as shown in Fig.4a,b. Unlike its variants, FilmCPI exhibits minimal optimization for the compound encoder but the greatest optimization for the protein encoder. This observation provides empirical support for our second argument and the correct choice of ECFPs for compound features. To reveal the optimization imbalance, we examine the differences among FilmCPI, FilmCPI_pc_, and FilmCPI_concat_, by discussing the error back-propagation process during training. Mathematical details for interpretation^26^ are displayed in Supplementary Section 2. Intriguingly, this observation challenges the traditional design concepts in previous methods, ^5,6^ which incorporate richer molecular features or better fusion methods for training to enhance their capacities.

## Discussion

Previous CPI prediction models^2,4–6,8,27^ implicitly assume that utilizing richer features would result in improved performance. However, this idea fails to account for the interdependent nature of the two distinct modalities involved in CPI predictors during training. The challenge of optimization imbalance^28,29^ has also been observed in those typical multimodal models. Actually, the concepts manifest differently in the compound-protein system compared to the typical multimodal system. For CPI prediction, the protein modality and the compound modality together define the label to be predicted. Contrarily, all modalities for a given pair point to the same label in a normal multimodal system. Besides, the solution to address the optimization imbalance in a typical multimodal system is to balance the learning rates for different modalities. However, this solution differs from the optimization dynamics of FilmCPI observed in Fig.4.

**Figure 4.**
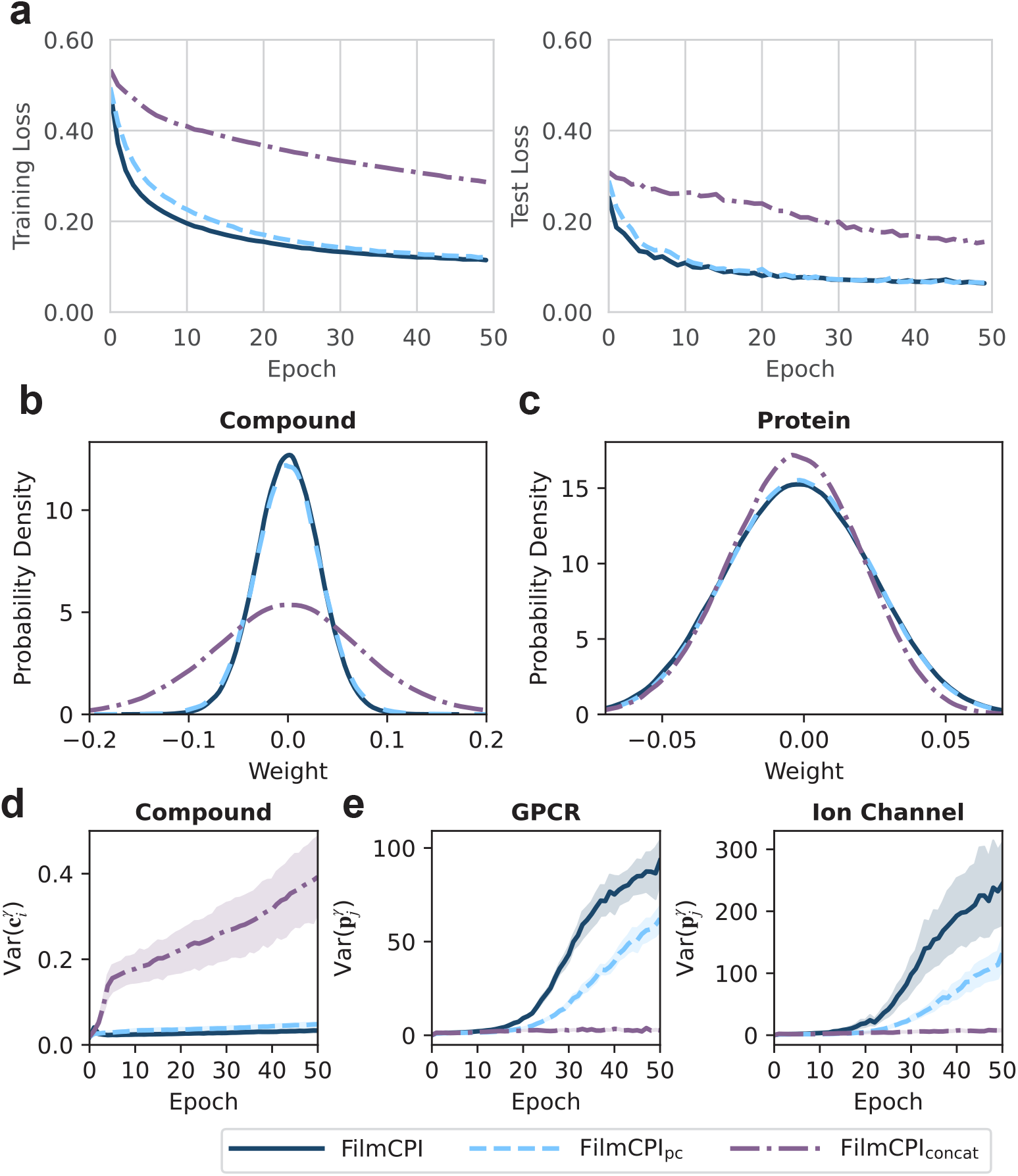
Analysis of optimization dynamics. **a**, Training loss and test loss of FilmCPI and its variants. **b**, Weight distributions in the compound encoders of FilmCPI and its variants. **c**, Weight distributions in the protein encoders of FilmCPI and its variants. **d**, Evolution of the variance of the compound representations within FilmCPI and its variants. **e**, Evolution of the variance of two protein representations (unseen GPCRs (left panel) & seen ion channels (right panel)) within FilmCPI and its variants.

This work primarily highlights the importance of the overlooked data imbalance. To this end, FilmCPI only employs ECFPs and protein sequences as inputs, making it to be the simplest yet effective CPI prediction model to our best knowledge. The simplicity of FilmCPI enables us to clearly establish a connection between the data imbalance observed in compound-protein datasets and the optimization imbalance inherent in models. This finding provides valuable insights for the design of both sequence-based and structure-based methods. For practical usage, FilmCPI is an ideal drug discovery tool for resource-constrained labs or can be utilized for preliminary screening before employing high-accuracy models such as AlphaFold3.^30^ On the other hand, the simplicity of FilmCPI suggests that there are numerous avenues for potential enhancements to this model. When considering the compound features, the utilization of ECFPs may be considered relatively basic. Recently, graph neural networks have become popular in the field of molecular representation learning. ^31,32^ Incorporating our above analysis, leveraging the more nuanced molecular representations of pre-trained molecular models^33–35^ may be a more appropriate choice. Besides, FilmCPI serves as a vanilla model in this work, which can incorporate some training strategies for further enhancements. For instance, a learning-to-rank strategy^36^ can be employed to learn the appropriate ranking orders, potentially even utilizing data from diverse evaluation metrics. Domain adaption^37^ and meta-learning^38^ approaches are valuable tools for improving the capacity of FilmCPI to handle some small protein families.

Compared to sequence-based CPI data, the amount of available structure-based CPI is limited. For instance, the number of protein-ligand co-complex structures in the latest PDBBind dataset^39^ does not exceed 20,000. While AlphaFold2 is a practical tool for predicting protein structures, for the structural models of the human proteome predicted by it, only 58% of the residues are modeled with high confidence.^40^ Although AlphaFold2 demonstrates excellent performance in predicting the overall backbone structure, it often lacks sufficient accuracy in details for structure-based drug discovery.^22^ In this context, the strengths of FilmCPI become evident. FilmCPI can leverage more CPI data compared to these structure-based methods. Furthermore, the scalability of FilmCPI illustrated in Fig.3a, along with the transferability exhibited in Fig.3b highlights its ability to handle large-scale data and further leverage such data to aid scenarios with limited data.

## Methods

### FilmCPI architecture

#### Features and corresponding encoders

Fig.1a exhibits the features and their corresponding encoders in FilmCPI for any pair {**c**_*i*_, **p**_*j*_}. For any compound **c**_*i*_, we first utilize RDKit,^41^ an open-source software, to change its SMILES into ECFP with diameter equal to 4 (ECFP4), as the compound input 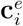. Then, we apply a fully connected (FC) layer to obtain the compound representations 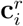, depicted in Eq.1.

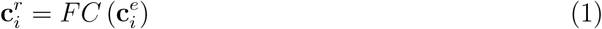

For any protein **p**_*j*_, we employ its amino acids sequences 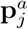 as inputs. For the protein module, we employ two 1D convolutional neural networks (CNNs) and a global max pooling layer to encode 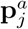 into protein representation 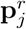, illustrated in Eq.2.

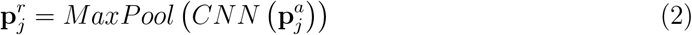

The first layer is a standard convolutional layer, while the second is a dilated convolutional layer.^42^ The dilated convolutional neural network is designed to expand the receptive field without increasing trainable parameters. The receptive field *F* is defined by Eq.3, where *k* is the kernel size and *r* is the dilation rate. From Eq.3, if the dilation rate *r* is set to 1, the receptive field is equal to the kernel size, meaning this convolutional layer is a standard convolutional layer. Conversely, if the dilation rate *r* is larger than 1, the receptive field becomes larger than the kernel size, indicating that this convolutional layer is a dilated convolutional layer.

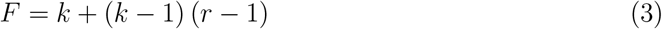

We employ Softplus^43^ as the activation function in this work to obtain more features since it has non-zero outputs. The details are illustrated in Eq.4, where *x* denotes any input.

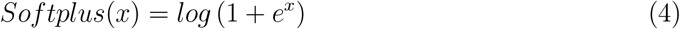

### Feature-wise linear modulation

Feature-wise linear modulation (Film)^19^ is a conditional normalization method. Similar techniques have been widely and successfully used for image stylization^44^ and visual question answering.^45^ That’s why we attempt to adopt the Film module to scale and shift 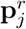 with 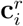. To be specific, 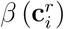 and 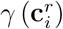 are learned from the compound representation 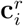 through **B** and **G** respectively, shown in Eq.5 and Eq.6. Then, they are used to modulate the protein representation 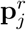, and the output of the Film module **X**_*i,j*_ is shown in Eq.7.

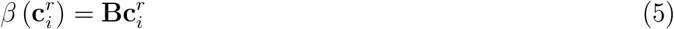

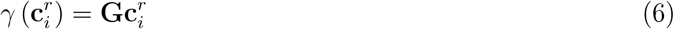

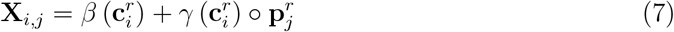

### Experimental settings

#### Datasets

Since different baselines in this work adopt diverse compound features, we aim to investigate the impact of dataset size, which is proportional to the number of compounds, to validate the robustness of FilmCPI. Thus, we collect two datasets with different sizes from the original BindingDB^15^ dataset with different thresholds for negative data. We first subset a smaller dataset named BindingDB_s_ considering a compound-protein pair to be positive only if its IC50 is less than 100 nM and negative only if its IC50 was greater than 10,000 nM.^5,21^ Next, we obtain BindingDB_b_ with IC50 greater than 1,000 nM to increase the amount of interaction data. We exclude all CPI pairs where the compounds only have one type of pair (positive or negative). Besides, we conduct experiments on BIOSNAP,^20^ the smallest dataset in this study. To further assess the practical values of these sequence-based methods, we select interaction data from two significant membrane protein families, ^22^ GPCR and ion channel, within BindingDB_b_ as the test set. The remaining data is used for training and validation purposes. Notably, the number of positive samples is larger than that of the negative samples in the GPCR dataset, which is in line with BindingDB_b_. However, the positive-to-negative sample ratio in the ion channel dataset is contrary, making the experiments more convincing. Statistics of datasets can be found in Supplementary Table 1,2.

#### Baselines and FilmCPI variants

To demonstrate the effectiveness of FilmCPI, we compare it against three representative models: DeepDTA, ^2^ MolTrans,^6^ and DrugBAN.^5^ These models differ significantly in their compound features and architectures. DeepDTA utilizes two convolutional modules to separately encode SMILES and protein sequences. MolTrans employs two transformer^46^ modules to encode sub-structural SMILES and protein sequences, respectively, followed by a convolutional module for interaction learning. DrugBAN adopts graph networks and convolutional networks to encode molecular graphs and protein sequences followed by a bilinear attention network.^47^ For a fair comparison, we adopt the recommended model hyperparameter settings described in their original papers.

We also include two variants of FilmCPI noted FilmCPI_pc_ and FilmCPI_concat_ as baselines to determine the values of FilmCPI design. Concretely, FilmCPI_pc_ exchanges the position of **c**_*i*_ and **p**_*j*_ in Eq.7, while FilmCPI_concat_ replaces the Film module with the concatenation operation.

#### Details of implementation

FilmCPI is implemented in Python 3.8 and PyTorch^48^ 1.13.1. The length of ECFP4 is set to 1,024, and the hidden dimension of the compound feature encoder is set to 256. The maximum sequence length of proteins is configured to 1,024. The protein feature encoder comprises two 1D CNN layers with the number of filters [256, 256], kernel sizes [3, 9], and dilation rates [1, 2].

When training FilmCPI, we set the batch size to 256 and employ Adam^49^ optimizer with a learning rate of 10^−4^. For both traditional evaluation and cross-family evaluation, we train FilmCPI for 50 epochs. Besides, we split these datasets using five different random seeds ranging from 0 to 4 and employ AUROC and AUPRC as the metrics to evaluate the models. The final result is the average to ensure the reliability of our findings. When discussing the optimization dynamics, we train FilmCPI and its variants in BindingDB_b_ without GPCRs for 50 epochs. All experiments are conducted in an NVIDIA A6000.

## Code and Data availability

The codes and data of this study will be added in https://github.com/Austin13579/fil_mcpi after acceptance.

## Acknowledgements

This work is supported by a start-up grant (grant no.: 9610591) for New Faculty and an internal grant (grant no.: 7006055) from the City University of Hong Kong to C.C.A.F. and the general grant (project no.: JCYJ20230807115001004) from the Science, Technology and Innovation Commission of Shenzhen Municipality to C.C.A.F. (the Shenzhen Research Institute, City University of Hong Kong).

