## Supplemental Information for "Utilizing data imbalance to enhance compound-protein interaction prediction models"

### Supplementary Method for "Utilizing data imbalance to enhance compound-protein interaction prediction models"

#### 1. Statistics of datasets

The number of proteins, compounds, interactions, positive pairs and negative pairs in three datasets, namely $\mathrm{BindingDB_b}$, $\mathrm{BindingDB_s}$, and BIOSNAP, are shown in Table1. The number of proteins, compounds, interactions, positive pairs and negative pairs in GPCR and ion channel dataset from $\mathrm{BindingDB_b}$ are shown in Table2.

Table 1: Details of $\mathrm{BindingDB_b}$, $\mathrm{BindingDB_s}$, and BIOSNAP.

| Datasets | #Proteins | #Compounds | #Interactions | # Positive | # Negative |
|---|---|---|---|---|---|
| $\mathrm{BindingDB_b}$ | 4005 | 50020 | 213143 | 115774 | 97369 |
| $\mathrm{BindingDB_s}$ | 3085 | 16854 | 76197 | 45939 | 30258 |
| BIOSNAP | 2181 | 4505 | 27464 | 13830 | 13634 |

Table 2: Details of GPCR and ion channel from $\mathrm{BindingDB_b}$.

| Protein family | #Proteins | #Compounds | #Interactions | # Positive | # Negative |
|---|---|---|---|---|---|
| GPCR | 162 | 5973 | 14235 | 8695 | 5540 |
| Ion channel | 41 | 4317 | 7974 | 2115 | 5859 |

### 2. Mathematical analysis

In this work, we aim to briefly illustrate the optimization mechanisms of FilmCPI and its variants from a mathematical perspective. Actually, the compound-protein interaction system is a multi-modal system since it contains two kinds of inputs. In the following part, we first follow the previous work to analyze the optimization dynamics of the concatenation operation,[1] thereby discussing the role of feature-wise linear modulation[2] (Film) module.

For any compound $\mathbf{c}_i$, we convert Simplified Molecular-Input Line Entry System (SMILES)[3] into its corresponding extended-connectivity fingerprint (ECFP)[4] $\mathbf{c}_i^e$ in this work. For any protein $\mathbf{p}_j$, we utilize its amino acids sequences $\mathbf{p}_j^a$. Then, protein representation $\mathbf{p}_j^r$ and compound representations $\mathbf{c}_i^r$ are integrated to $\mathbf{X}_{i,j}$ before going through the decoder for final prediction. In this study, $\mathbf{W}^c$, $\mathbf{W}^p$ and $\mathbf{W}^D$ are the weight matrices for the compound encoder, the protein encoder and the decoder. Besides, we set $L$ as the loss function, and $\boldsymbol{\delta}^D$ is the error signal fed from the decoder.

#### 2.1 Different optimization speeds

First of all, we discuss the concatenation operation. In this situation, we have

$$\mathbf{X}_{i,j} = \text{concatenate}\left(\mathbf{c}_i^r, \mathbf{p}_j^r\right), \tag{1}$$

where we abstractly formulate the two encoders with:

$$\mathbf{c}_i^r = \varphi\left(\mathbf{W}^c, \mathbf{c}_i^e\right) \tag{2}$$

$$\mathbf{p}_j^r = \varphi\left(\mathbf{W}^p, \mathbf{p}_j^a\right) \tag{3}$$

For the optimization of the protein encoder and the compound encoder, we have

$$\frac{\partial L}{\partial \mathbf{W}^p} = \left\{\left[\left(\mathbf{W}^D\big|_p\right)^T \boldsymbol{\delta}^D\right] \circ \frac{\partial \varphi}{\partial \mathbf{W}^p}\left(\mathbf{W}^p, \mathbf{p}_j^a\right)\right\} \tag{4}$$

$$\frac{\partial L}{\partial \mathbf{W}^c} = \left\{\left[\left(\mathbf{W}^D\big|_c\right)^T \boldsymbol{\delta}^D\right] \circ \frac{\partial \varphi}{\partial \mathbf{W}^c}\left(\mathbf{W}^c, \mathbf{c}_i^e\right)\right\}. \tag{5}$$

Based on Eq.2 and Eq.3, the gradients of loss function $L$ for both the protein encoder $\mathbf{W}^p$ and the compound encoder $\mathbf{W}^c$ are influenced by their respective inputs, namely $\mathbf{p}_j^a$ and $\mathbf{c}_i^e$, assuming the influence of the decoder is ignored. Since proteins are larger in size but fewer in number compared to compounds in most datasets, the optimization of the protein encoder should be more challenging than that of the compound encoder if utilizing a concatenation operation. This aligns with the observations from previous work.[5] Besides, awful performances of $\text{FilmCPI}_{\text{concat}}$, exhibited in the main text, support our argument that different optimization speeds are crucial factors for different encoders during training.

#### 2.2 Optimization dynamics of FilmCPI

If we use the compound representations $\mathbf{c}_i^r$ to conditionally normalize the protein representations $\mathbf{p}_j^r$ with Film module, the output of FilmCPI is defined by

$$\mathbf{X}_{i,j} = \beta\left(\mathbf{c}_i^r\right) + \gamma\left(\mathbf{c}_i^r\right) \circ \mathbf{p}_j^r, \tag{6}$$

where

$$\mathbf{c}_i^r = \varphi\left(\mathbf{W}^c, \mathbf{c}_i^e\right) \tag{7}$$

$$\mathbf{p}_j^r = \varphi\left(\mathbf{W}^p, \mathbf{p}_j^a\right) \tag{8}$$

and **B** and **G** are weight matrices with the same dimensions.

Then, the gradient descents for **B** and **G** are

$$\mathbf{B}\left(t+1\right) = \mathbf{B}\left(t\right) - \lambda \frac{\partial L}{\partial \mathbf{B}}\left(t\right) \tag{9}$$

$$\mathbf{G}\left(t+1\right) = \mathbf{G}\left(t\right) - \lambda \frac{\partial L}{\partial \mathbf{G}}\left(t\right) \tag{10}$$

where

$$\frac{\partial L}{\partial \mathbf{G}} = \left(\nabla_{\mathbf{X}} L \circ \mathbf{p}_j^r\right) \otimes \mathbf{c}_i^r \tag{11}$$

$$\frac{\partial L}{\partial \mathbf{B}} = \nabla_{\mathbf{X}} L \otimes \mathbf{c}_i^r \tag{12}$$

Therefore, for the compound encoder, we have

$$\frac{\partial L}{\partial \mathbf{W}^c} = \left\{\left[\mathbf{B}^T \left(\mathbf{W}^D\right)^T \boldsymbol{\delta}^D + \mathbf{G}^T \left(\left(\left(\mathbf{W}^D\right)^T \boldsymbol{\delta}^D\right) \circ \mathbf{p}_j^r\right)\right] \circ \frac{\partial \sigma}{\partial \mathbf{W}^c}\left(\mathbf{W}^c, \mathbf{c}_i^e\right)\right\} \tag{13}$$

For the protein encoder, we have

$$\frac{\partial L}{\partial \mathbf{W}^p} = \left\{\left[\left(\mathbf{W}^D\right)^T \boldsymbol{\delta}^D\right] \circ \left(\mathbf{G}\mathbf{c}_i^r\right) \circ \frac{\partial \sigma}{\partial \mathbf{W}^p}\left(\mathbf{W}^p, \mathbf{p}_j^a\right)\right\} \tag{14}$$

#### 2.3 Optimization dynamics of $\mathbf{FilmCPI_{pc}}$

If we exchange the positions of the compound representations in the Film module, the output of $\text{FilmCPI}_{\text{pc}}$ is defined by

$$\mathbf{X}_{i,j} = \beta\left(\mathbf{p}_j^r\right) + \gamma\left(\mathbf{p}_j^r\right) \circ \mathbf{c}_i^r, \tag{15}$$

where

$$\beta\left(\mathbf{p}_j^r\right) = \mathbf{B}\mathbf{p}_j^r \tag{16}$$

$$\gamma\left(\mathbf{p}_j^r\right) = \mathbf{G}\mathbf{p}_j^r \tag{17}$$

$$\frac{\partial L}{\partial \mathbf{G}} = \left(\nabla_{\mathbf{X}} L \circ \mathbf{c}_i^r\right) \otimes \mathbf{p}_j^r \tag{18}$$

$$\frac{\partial L}{\partial \mathbf{B}} = \nabla_{\mathbf{X}} L \otimes \mathbf{p}_j^r \tag{19}$$

For the protein encoder, we have

$$\frac{\partial L}{\partial \mathbf{W}^p} = \left\{\left[\mathbf{B}^T \left(\mathbf{W}^D\right)^T \boldsymbol{\delta}^D + \mathbf{G}^T \left(\left(\left(\mathbf{W}^D\right)^T \boldsymbol{\delta}^D\right) \circ \mathbf{c}_i^r\right)\right] \circ \frac{\partial \sigma}{\partial \mathbf{W}^p}\left(\mathbf{W}^p, \mathbf{p}_j^a\right)\right\} \tag{20}$$

For the compound encoder, we have

$$\frac{\partial L}{\partial \mathbf{W}^c} = \left\{\left[\left(\mathbf{W}^D\right)^T \boldsymbol{\delta}^D\right] \circ \left(\mathbf{G}\mathbf{p}_j^r\right) \circ \frac{\partial \sigma}{\partial \mathbf{W}^c}\left(\mathbf{W}^c, \mathbf{c}_i^e\right)\right\} \tag{21}$$

#### 2.4 Comparison of optimization dynamics between FilmCPI and $\text{FilmCPI}_{\text{pc}}$

For the optimization of FilmCPI, the parameters $\mathbf{G}$, $\mathbf{W}^c$ and $\mathbf{W}^p$, depicted in Eq.11,13,14, are influenced by both compounds and proteins. Similarly, for the optimization of $\text{FilmCPI}_{\text{pc}}$, $\mathbf{G}$, $\mathbf{W}^c$ and $\mathbf{W}^p$, depicted in Eq.18,20,21, are also affected by both compounds and proteins. This interdependent optimization aims to better integrate the two modalities. Since the optimization is difficult to the protein encoders, the optimization to compound encoders is constrained in both FilmCPI and $\text{FilmCPI}_{\text{pc}}$. As a result, both FilmCPI and $\text{FilmCPI}_{\text{pc}}$ outperform $\text{FilmCPI}_{\text{concat}}$ in random split and unseen compound split.

The key difference between the two settings lies in $\mathbf{B}$. As shown in Eq.12, $\mathbf{B}$ in FilmCPI is influenced solely by compounds, whereas in $\text{FilmCPI}_{\text{pc}}$, $\mathbf{B}$ is influenced only by proteins, shown in Eq.19. This indicates that $\text{FilmCPI}_{\text{pc}}$ is more dominated by protein representations compared to FilmCPI. However, since optimizing the protein encoder is more difficult than optimizing the compound encoder based on the analysis of $\text{FilmCPI}_{\text{concat}}$, the optimization of the protein encoder in $\text{FilmCPI}_{\text{pc}}$ is less effective than that in FilmCPI. With the expansion of datasets, this difference results in progressively poorer performances on unseen proteins in $\text{FilmCPI}_{\text{pc}}$, leading to progressively better outcomes on unseen proteins in FilmCPI.

### 3. Numerical results

Table 3 exhibits the results of all models in the unseen protein split, in $\text{BindingDB}_{\text{b}}$, $\text{BindingDB}_{\text{s}}$, and BIOSNAP, respectively. Table 4 shows the performance of models evaluated with G-protein-coupled receptors (GPCRs) and ion channels.

Table 3: Performance (average AUROC and AUPRC over five random runs) comparison in $BindingDB_b$, $BindingDB_s$ and BIOSNAP with the unseen protein split.

| Dataset | $BindingDB_b$ | | $BindingDB_s$ | | BIOSNAP | |
|---|---|---|---|---|---|---|
| Metric | AUROC | AUPRC | AUROC | AUPRC | AUROC | AUPRC |
| DeepDTA | 0.619±0.040 | 0.629±0.067 | 0.652±0.043 | 0.748±0.049 | 0.800±0.027 | 0.836±0.029 |
| MolTrans | 0.537±0.020 | 0.573±0.034 | 0.563±0.046 | 0.686±0.033 | 0.691±0.021 | 0.721±0.034 |
| DrugBAN | 0.592±0.035 | 0.611±0.044 | 0.601±0.041 | 0.713±0.040 | 0.668±0.033 | 0.705±0.059 |
| $FilmCPI_{pc}$ | 0.549±0.036 | 0.571±0.070 | 0.603±0.048 | 0.707±0.055 | 0.781±0.020 | 0.818±0.029 |
| $FilmCPI_{concat}$ | 0.627±0.053 | 0.647±0.063 | 0.633±0.025 | 0.736±0.033 | 0.769±0.025 | 0.798±0.033 |
| FilmCPI | **0.668±0.036** | **0.695±0.040** | **0.694±0.043** | **0.788±0.044** | **0.816±0.013** | **0.851±0.018** |

Table 4: Performance (average AUROC and AUPRC over five random runs) comparison in GPCR or ion channel from $BindingDB_b$.

| Split Type | GPCR | | Ion Channel | |
|---|---|---|---|---|
| Metric | AUROC | AUPRC | AUROC | AUPRC |
| DeepDTA | 0.603±0.010 | 0.697±0.008 | 0.543±0.147 | 0.331±0.126 |
| MolTrans | 0.506±0.013 | 0.623±0.014 | **0.673±0.070** | 0.444±0.100 |
| DrugBAN | 0.580±0.017 | 0.670±0.016 | 0.520±0.041 | 0.282±0.028 |
| FilmCPI | **0.614±0.015** | **0.713±0.015** | 0.647±0.042 | **0.457±0.052** |

(4) Rogers, D.; Hahn, M. Extended-connectivity fingerprints. *Journal of chemical informa-*

*tion and modeling* **2010**, *50*, 742–754.

(5) Chen, L.; Tan, X.; Wang, D.; Zhong, F.; Liu, X.; Yang, T.; Luo, X.; Chen, K.; Jiang, H.; Zheng, M. TransformerCPI: improving compound–protein interaction prediction by sequence-based deep learning with self-attention mechanism and label reversal experiments. *Bioinformatics* **2020**, *36*, 4406–4414.
